# Gene expression is stable despite widespread cis and trans regulatory divergence in *Saccharomyces* yeasts

**DOI:** 10.64898/2026.06.03.729905

**Authors:** Danithza S. Rojas, Gregory I. Lang

## Abstract

Regulatory evolution can alter phenotypes, but cis- and trans-regulatory mechanisms may also diverge extensively while total transcript abundance remains stable. Comparisons of parental expression with allele-specific expression in F1 hybrids provide a framework for separating cis- and trans-regulatory effects because both parental alleles are measured in a shared trans-regulatory environment. Here, we analyzed RNA sequencing data from *Saccharomyces cerevisiae*, *Saccharomyces paradoxus*, and their F1 hybrid. Regulatory divergence was widespread, with 61.3% of tested orthologs showing significant divergence in at least one cis or trans component. However, hybrid expression remained largely conserved, with 81.6% of genes not significantly different from either parent. Compensatory cis-trans divergence predominated over reinforcing divergence, consistent with widespread buffering of transcript abundance. To connect genome-wide patterns to mechanism, we analyzed the strongly cis-diverged locus *LYS2* and found species differences in promoter architecture, including an *S. cerevisiae*-specific AT-rich insertion, altered spacing among candidate regulatory features, and a promoter-proximal TATA-like element unique to *S. cerevisiae*. Sequence-based nucleosome prediction suggests that these differences create a broader promoter-proximal nucleosome-depleted region in *S. cerevisiae* than in *S. paradoxus*. We also quantified allele-resolved intron retention and found that splicing was broadly conserved, with only rare locus-specific hybrid-associated shifts. Together, these results show that regulatory divergence is widespread but often buffered in the hybrid, whereas post-transcriptional divergence is comparatively limited.

## INTRODUCTION

Gene expression levels can remain stable even when the regulatory mechanisms that produce them have diverged. This distinction is central to understanding regulatory evolution. Genetic variation can affect gene expression in one of two general ways. Cis-regulatory elements influence expression locally and are physically linked to the gene they regulate. Trans-acting factors are diffusible regulators that can be encoded anywhere in the genome and can regulate any number of genes. Variation in cis-regulatory or trans-acting factors may alter the architecture of gene regulation without necessarily producing large changes in total transcript abundance. In some cases, opposing cis and trans effects can produce compensatory regulatory divergence, allowing transcript output to remain stable even as the underlying regulatory mechanisms diverge. Such hidden regulatory divergence is important because it can reveal how transcript output is maintained over evolutionary time while the underlying cis and trans components continue to change. Therefore, distinguishing between local cis-regulatory effects and diffusible trans-regulatory effects is essential for understanding not only how transcriptomes diverge, but also how gene expression stability is preserved despite regulatory evolution (King and Wilson, 1975; Carroll, 2005; Wray, 2007; Wittkopp and Kalay, 2012; Hill et al., 2021; Signor and Nuzhdin, 2018).

The use of hybrids between closely related species provides a framework for separating the contributions of cis- and trans-regulatory elements because the two parental alleles coexist within the same nucleus and are exposed to the same pool of diffusible trans-acting factors. In this shared trans-regulatory environment, allele-specific expression differences between the two parental alleles are interpreted as evidence of cis-regulatory divergence (Cowles et al., 2002). In contrast, trans effects are inferred by comparing total expression differences between the parental species with the allele-specific expression pattern observed in the hybrid (Wittkopp et al., 2004; Tirosh et al., 2009; Schaefke et al., 2013). Research across diverse systems, including *Saccharomyces* yeasts, Drosophila, and mammals, has shown that both cis and trans changes are common and often accumulate in opposing directions, a pattern consistent with compensatory evolution under stabilizing selection on transcript abundance (Landry et al., 2005; McManus et al., 2010; Metzger et al., 2017; Hill et al., 2021). Thus, even when the underlying regulatory mechanisms have diverged substantially, overall mRNA levels can remain relatively stable.

These observations are especially relevant in hybrids, where regulatory components that evolved in different parental lineages must function together in a shared cellular environment. Co-evolved cis and trans changes within each species can stabilize transcript output in the hybrid and keep expression within the parental range, whereas hybridization can also uncover regulatory mismatches that are not apparent from parental expression alone (Landry et al. 2005; Tirosh et al. 2009; Herbst et al. 2017). The *Saccharomyces cerevisiae/Saccharomyces paradoxus* system provides a useful model for examining this balance. These closely related yeasts remain cross-compatible while harboring substantial regulatory divergence. Previous studies in *Saccharomyces* have documented genome-wide cis and trans differences, dominance of many trans effects, complex trans-regulatory architectures, and divergence in expression dynamics across environments (Tirosh et al. 2009; Schaefke et al. 2013; Metzger et al. 2017; Albert et al. 2018; Krieger et al. 2020; Shih and Fay 2021).

Here, we combine cis-trans regulatory decomposition with analysis of total expression in the F1 hybrid. Rather than asking only whether genes differ in expression between *S. cerevisiae* and *S. paradoxus*, we ask how those differences are encoded, including as cis, trans, compensatory, or reinforcing effects, and whether expression in the hybrid remains within the parental range or reveals misregulation. To address these questions, we integrate total RNA sequencing, allele-specific expression, and allele-resolved intron-retention analysis in *Saccharomyces cerevisiae*, *Saccharomyces paradoxus*, and their F1 hybrid. This approach allows us to ask how regulatory divergence is partitioned into cis and trans components, how these architectures shape hybrid expression patterns, and whether promoter architecture or post-transcriptional regulation reveals additional layers of divergence. We find widespread compensatory cis-trans evolution together with generally stable hybrid expression, rare and locus-specific hybrid-associated shifts in intron retention, and a candidate cis-regulatory mechanism at *LYS2* that is consistent with divergent promoter architecture and predicted chromatin organization.

## MATERIALS AND METHODS

### Yeast strains and media

We analyzed gene expression and splicing in diploid *Saccharomyces cerevisiae* (*S.c*) and *S. paradoxus* (*S.p*) parents and in their F1 heterozygous diploid hybrid. All cultures were grown in rich medium (YPD; 1% yeast extract, 2% peptone, 2% glucose) at 30 °C with shaking and harvested at mid-log phase (OD600 ≈ 0.6). We collected 2–5 independent biological replicates per genotype. Cells were pelleted by rapid centrifugation, flash-frozen in liquid nitrogen, and stored at −80 °C before RNA extraction.

### RNA extraction and library preparation

Total RNA was extracted using hot acid phenol-chloroform followed by silica-column cleanup (Qiagen RNeasy). RNA quantity and integrity were assessed by Qubit fluorometry and Bioanalyzer analysis, and only samples with RIN ≥ 7 were used for library preparation. Poly(A)+ RNA was enriched from 1–2 µg total RNA using the NEBNext Poly(A) mRNA Magnetic Isolation Module (NEB E7490). Strand-specific RNA-seq libraries were prepared with the NEBNext Ultra II Directional RNA Library Prep Kit (NEB E7760) according to the manufacturer’s dUTP-based protocol. Indexed libraries were amplified using 10 PCR cycles, size-selected with AMPure beads, and quantified by Qubit and Bioanalyzer to confirm appropriate insert size distributions. Libraries were pooled and sequenced on an Illumina NovaSeq platform to generate paired-end reads of at least 75 bp. We recovered between 10 and 20 million read pairs per sample.

### Read processing and alignment

Adapters and low-quality bases were removed with Trim Galore/Cutadapt (Martin, 2011); read quality was summarized across samples with MultiQC (Ewels et al., 2016). We constructed a combined reference genome by concatenating the *S. c* and *S. p* assemblies and prefixing chromosome names to distinguish species while preserving chromosome numbering. The corresponding gene-annotation (GTF) files were merged after harmonizing one-to-one ortholog identifiers. Trimmed reads were aligned to this combined hybrid genome with STAR in two-pass, splice-aware mode (Dobin et al., 2013). Alignment outputs were coordinate-sorted and PCR duplicates were marked. These BAM files were used for all downstream analyses, including gene-level quantification, allele-specific expression, and intron-retention profiling.

### Gene quantification and filtering

Gene-level counts were obtained with featureCounts from the Subread package using the merged hybrid GTF and requiring uniquely mapped, properly paired reads (Liao et al., 2014). One-to-one orthologs were assigned a shared gene identifier across species. To avoid testing genes with negligible expression, we retain genes with at least 1 count per million (CPM) in at least two samples. After filtering, 5,243 genes remained for analysis from a starting set of approximately 5,400 orthologs. All downstream analyses were performed in R.

### Differential expression between parents

Differential expression between the *S. c* and *S. p* diploid parents was tested with DESeq2 (Love et al., 2014). Size factors were estimated using the median-of-ratios method, and gene-wise dispersions were fit and shrunk with the package’s empirical Bayes framework. Wald tests were used for inference, and P values were adjusted for multiple testing by the Benjamini-Hochberg false discovery rate (FDR) procedure. Unless otherwise noted, genes were considered significantly divergent at padj < 0.005 and absolute log2 fold-change (|LFC|) ≥ 0.58, corresponding to an approximately 1.5-fold difference. In figures where low-count genes required more stable effect-size estimates, we used apeglm shrinkage for LFC visualization without changing significance calls (Zhu et al., 2019). For interpretation, we recorded the DESeq2 baseMean for each gene and calculated group-specific mean expression values for *S. c*, *S. p*, and the hybrid.

### Hybrid expression inheritance classification

To classify hybrid expression relative to the parents, we modeled the two diploid parents and the hybrid jointly in DESeq2 and extracted the contrasts Hybrid vs. *S. c* and Hybrid vs. *S. p*, again using LFC shrinkage for stability. We then assigned each gene to an inheritance category using a scheme that required both statistical significance and logical placement relative to parental means.

Nondifferential genes were those for which hybrid mean expression lay within the parental range and did not differ significantly from either parent. Additive genes were those for which the hybrid mean lay strictly between the two parents and differed significantly from both, with opposite-sign LFCs relative to each parent. Dominant genes were those for which the hybrid did not differ significantly from one parent but did differ significantly from the other, consistent with expression matching one parental state. Overdominant or underdominant (transgressive) genes were those for which hybrid expression was significantly above both parents or below both parents, respectively, with padj < 0.005 and |LFC| ≥ 0.58 in both hybrid-versus-parent contrasts.

If a hybrid mean fell outside the parental range but failed the significance criteria against one or both parents, we labeled the gene as outside but not significant rather than calling it transgressive. This avoided overcalling overdominance or underdominance and ensured that only robust cases of hybrid misexpression were placed in transgressive classes.

### Allele-specific expression and cis–trans decomposition

Allele-specific expression (ASE) in the hybrid was used to estimate cis-regulatory divergence. To minimize mapping bias, we used a WASP-based filtering strategy during alignment (van de Geijn et al., 2015), an approach consistent with published best practices for allelic-expression analysis (Castel et al., 2015). For each read overlapping a known *S. cerevisiae/S. paradoxus* single-nucleotide polymorphism (SNP), the overlapping allele was computationally swapped and the read was remapped; reads that aligned only in their original form were discarded. This procedure yielded bias-reduced BAM files for ASE quantification. High-quality allele-specific counts per SNP were then obtained with GATK ASEReadCounter within the GATK framework (McKenna et al., 2010), requiring at least 20 reads per SNP and unique mapping to exonic sequence. Allelic counts were summed by gene and replicate.

For each gene in each replicate, deviation of *S. c* and *S. p* allele counts from the 50:50 null expectation was tested with a two-sided binomial test. We then converted each replicate result to a signed Z score based on the direction of imbalance [log2(*S. c* /*S. p*)] and combined replicate Z scores with Stouffer’s method. Resulting meta-analysis P values were adjusted for multiple testing (padjASE), and genes with padjASE < 0.005 and |log2 ASE| ≥ 0.58 were called as having significant cis-regulatory divergence. For each gene, the cis effect was defined as LFCASE = log2(*S. c* allele / *S. p* allele), together with its standard error. The trans component was estimated as LFCtrans = LFCparent − LFCASE. Standard errors for trans were obtained by error propagation, and trans P values were derived from Z tests followed by FDR correction (padjtrans).

Genes were assigned to regulatory classes using significance and effect-size thresholds for the cis and trans components. Genes with a significant cis effect (padjASE < 0.005 and |LFC| ≥ 0.58) but no significant trans effect were classified as “cis-only.” Genes with a significant trans effect (padjtrans < 0.005 and |LFC| ≥ 0.58) but no significant cis effect were classified as “trans-only.” Genes for which both cis and trans were significant and had the same sign were classified as “cis + trans, reinforcing,” whereas genes for which both components were significant but had opposite signs were classified as “cis + trans, compensatory.” Genes for which neither component passed the significance and effect-size criteria were classified as “conserved or ambiguous.” This category includes genes with no detectable regulatory divergence as well as borderline or low-effect cases that could not be resolved confidently. These criteria provided a high-confidence partitioning of regulatory divergence into mechanistic classes.

### Gene Ontology enrichment analysis

To test functional bias across regulatory classes, we performed Gene Ontology (GO) enrichment analysis focused on Biological Process and Molecular Function annotations using the Gene Ontology resource (Gene Ontology Consortium, 2019). For each regulatory class, the query set consisted of genes assigned to that class, and the background set consisted of all 5,243 tested genes. Enrichment was assessed with one-sided Fisher’s exact tests and corrected for multiple testing using the Benjamini-Hochberg procedure. Terms with FDR < 0.01 were considered significant, and we report representative enrichments together with their fold-enrichment values and adjusted P values. For follow-up analysis of the trans-only class, we further separated trans-only genes by hybrid inheritance mode into *S. p*-dominant and *S. c*-dominant subsets and repeated GO enrichment using the same 5,243 tested genes as the background.

### Intron retention quantification and allele-resolved comparisons

Intron retention (IR) was quantified with IRFinder v1.3 in directional mode (Middleton et al., 2017). IRFinder estimates an intron-level IR ratio by comparing reads that support intron retention with reads that support splicing.

We compared intron retention between each diploid parent and the corresponding allele in the F1 hybrid. The *S. c* diploid parent, with two biological replicates, was compared with the *S. c* allele in the hybrid, and the *S. p* diploid parent, with four biological replicates, was compared with the *S. p* allele in the hybrid. The hybrid had five biological replicates.

To focus on reliable intron-retention estimates, we retained introns with sufficient read support and consistent quantification across biological replicates. After filtering, 178 introns were analyzed in the *S. c* parent-to-hybrid allele comparison, and 149 introns were analyzed in the *S. p* parent-to-hybrid allele comparison.

Differential intron retention between each parent and the corresponding hybrid allele was tested using Fisher’s exact tests comparing retained and spliced read support, followed by Benjamini-Hochberg correction within each allele-specific comparison. Effect size was defined as ΔIR = IR hybrid allele minus IR parent. Differential intron-retention events were defined as those with absolute ΔIR greater than 0.1 and FDR less than 0.05. One additional event that narrowly missed the effect-size cutoff but showed strong statistical support was reported separately as a near-threshold event.

### *LYS2* promoter comparison and nucleosome prediction

To compare promoter architecture at the *LYS2* locus across *Saccharomyces* species, we identified the annotated *LYS2* ortholog in each species and extracted the 600 bp region upstream of the coding sequence. Promoter sequences were extracted from the corresponding genome FASTA files while accounting for gene orientation, so that all sequences were analyzed in the same transcriptional direction. The *S. c* and *S. p* promoters were included as the focal comparison, and three additional sensu stricto species, *S. mikatae* (*S. m*), *S. kudriavzevii* (*S. k*), and *S. uvarum* (*S. u*), were included to place these differences in a broader evolutionary context.

Promoter sequences were globally aligned and compared using three summary measures: percent identity across aligned non-gap positions, total insertion/deletion burden, and the number of separate insertion/deletion events. These measures were used to distinguish broad promoter sequence divergence from more localized architectural differences.

Because TATA boxes are sequence-variable, promoters were scanned for TATA-like elements using the degenerate consensus TATAWAWA, where W represents A or T. Overlapping matches were allowed, and motif positions were recorded relative to the coding-sequence start site so that negative values indicate upstream promoter positions. Motif scanning was performed with FIMO (Grant et al., 2011).

Intrinsic nucleosome occupancy was predicted for each promoter using NuPoP with the yeast model (Xi et al., 2010). NuPoP provides a predicted occupancy score at each promoter position, with higher values indicating greater predicted nucleosome occupancy. To compare promoter-proximal chromatin architecture across species, we summarized mean predicted occupancy in three regions of interest: an upstream indel-rich region near −390 bp, a broader core-promoter region from −150 to −50 bp, and a TATA-proximal region from −120 to −80 bp. These summaries were used to evaluate whether sequence differences at *LYS2* were associated with differences in predicted promoter-proximal nucleosome organization.

## RESULTS

### Regulatory divergence is pervasive and largely compensatory

To reveal the regulatory architecture underlying gene expression in *S. cerevisiae* and *S. paradoxus*, we measured gene expression in the diploid parents and the diploid F1 hybrid. Gene expression differences between species can arise from cis or trans regulatory effects. Cis effects can be estimated from allele-specific expression (ASE) in the hybrid, where both parental alleles are exposed to the same trans-regulatory environment (Fig. 1A). Trans effects can then be inferred by comparing ASE in the hybrid with total expression differences between the parental strains (Fig. 1A). Among 5,386 one-to-one orthologous genes, 5,243 genes (97.3%) satisfied expression-filtering criteria and were included in the cis-trans analysis. Using our combined effect-size and significance threshold of |log2 fold-change| ≥ 0.58 and adjusted p < 0.005, 3,212 genes (61.3%) showed significant regulatory divergence in at least one component, cis and/or trans.

**Figure 1.**
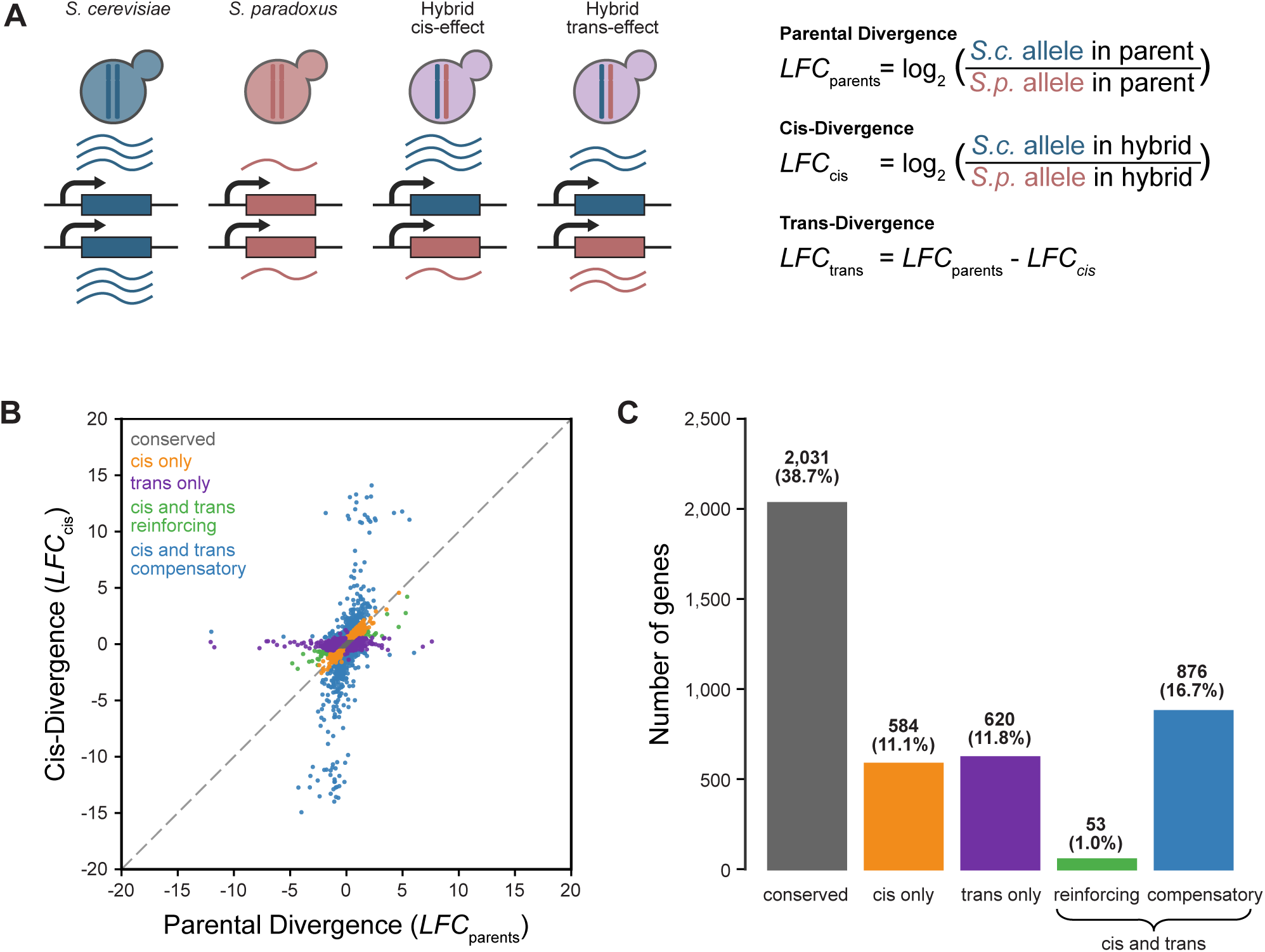
Experimental framework and genome-wide classification of cis- and trans-regulatory divergence in *S. cerevisiae × S. paradoxus.* **(A)** Conceptual schematic for partitioning regulatory divergence using parental expression and allele-specific expression (ASE) in the F1 hybrid. In the cis example, the two parental alleles differ in the hybrid, indicating allele-linked regulatory divergence. In the trans example, parental total expression differs whereas the two alleles are similarly expressed in the hybrid, indicating divergence in the shared trans environment. The equations used to calculate parental divergence, the cis component, and the inferred trans component are shown at right. *S. cerevisiae* is shown in blue, *S. paradoxus* in red, and the hybrid in purple. **(B)** Scatter plot of regulatory divergence across genes. The x-axis shows parental expression divergence and the y-axis shows the cis component measured by ASE in the hybrid. Points are colored by regulatory class: conserved, cis only, trans only, cis + trans reinforcing, and cis + trans compensatory. The dashed diagonal indicates equality between parental divergence and the cis component. **(C)** Number and percentage of genes assigned to each regulatory class.

Of the 3,212 divergent genes, 2,133 could be assigned to a resolved regulatory class (Fig. 1B and C). These included 584 cis-only genes, 620 trans-only genes, 876 compensatory cis + trans genes, and 53 reinforcing cis + trans genes, corresponding to approximately 11.1%, 11.8%, 16.7%, and 1.0% of all tested genes, respectively. The remaining 1,079 divergent genes showed significant but low-effect or ambiguous regulatory differences that did not meet the criteria for assignment to a specific cis-trans class. We therefore retained them in the genome-wide summary of regulatory divergence but did not include them in downstream class-specific comparisons that required a resolved regulatory architecture.

Among the resolved genes, nearly half (929 of 2,133) had both cis and trans contributions. Opposite-sign compensatory changes were much more common than same-direction reinforcing changes, with compensatory genes outnumbering reinforcing genes by approximately 16.5-fold. Together, these results show that regulatory divergence between *S.c* and *S.p* is widespread and is dominated by compensatory cis-trans architectures.

To determine whether genes in particular biological processes tend to share regulatory divergence, we performed Gene Ontology (GO) enrichment analysis using the 5,243 tested orthologs as the background set (Gene Ontology Consortium, 2019; Sup Fig. 1A–C). Significant enrichment was detected only in the conserved and trans-only classes, whereas the cis-only, compensatory, and reinforcing classes showed no significant biological process enrichment at the same threshold (Fig. 2A). This suggests that pathway-level organization is strongest among genes with conserved regulation and genes with trans-only divergence, while cis-containing classes are more functionally heterogeneous.

**Figure 2.**
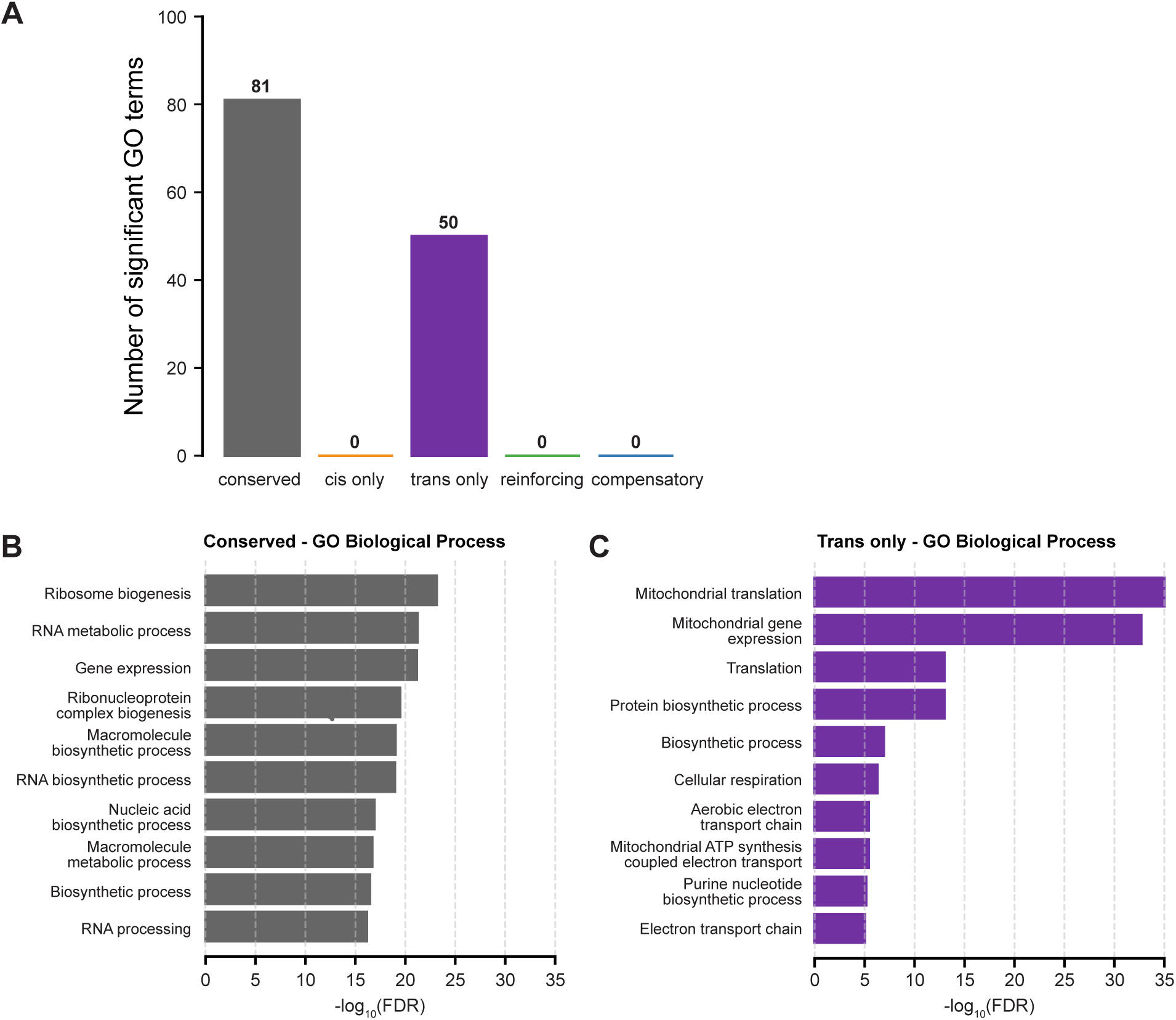
GO biological process enrichment reveals pathway-level organization concentrated in conserved and trans-only regulatory classes. **(A)** Number of significantly enriched GO biological process terms identified in each regulatory class using the same ortholog-restricted background of 5,243 tested genes. Enrichment was assessed for all classes under the same statistical framework, but only the conserved and trans-only classes yielded significant terms, with 81 and 50 enriched terms, respectively. The cis-only, cis + trans reinforcing, and cis + trans compensatory classes showed no significant biological process enrichment at the same threshold. **(B)** Top 10 enriched GO biological process terms for conserved genes, ranked from most to least significant. Conserved genes were enriched primarily for ribosome biogenesis, RNA metabolism, gene expression, ribonucleoprotein complex biogenesis, and related biosynthetic and RNA-processing functions, consistent with strong conservation of core housekeeping pathways. The terms shown are ribosome biogenesis (GO:0042254; FDR = 7.01 × 10^-24; n = 229), RNA metabolic process (GO:0016070; FDR = 5.95 × 10^-22; n = 597), gene expression (GO:0010467; FDR = 7.16 × 10^-22; n = 801), ribonucleoprotein complex biogenesis (GO:0022613; FDR = 3.37 × 10^-20; n = 253), macromolecule biosynthetic process (GO:0009059; FDR = 9.51 × 10^-20; n = 863), RNA biosynthetic process (GO:0032774; FDR = 1.12 × 10^-19; n = 530), nucleic acid biosynthetic process (GO:0141187; FDR = 1.23 × 10^-17; n = 540), macromolecule metabolic process (GO:0043170; FDR = 2.01 × 10^-17; n = 1070), biosynthetic process (GO:0009058; FDR = 3.43 × 10^-17; n = 1026), and RNA processing (GO:0006396; FDR = 6.97 × 10^-17; n = 286). **(C)** Top 10 enriched GO biological process terms for trans-only genes, ranked from most to least significant. Trans-only genes were enriched for mitochondrial translation, mitochondrial gene expression, cellular respiration, electron transport, and related energy-metabolism functions, indicating that trans-regulatory divergence is concentrated in coordinated mitochondrial and respiratory modules. The terms shown are mitochondrial translation (GO:0032543; FDR = 2.64 × 10^-37; n = 77), mitochondrial gene expression (GO:0140053; FDR = 2.04 × 10^-33; n = 80), translation (GO:0006412; FDR = 1.04 × 10^-13; n = 102), protein biosynthetic process (GO:0160307; FDR = 1.04 × 10^-13; n = 102), biosynthetic process (GO:0009058; FDR = 1.22 × 10^-7; n = 338), cellular respiration (GO:0045333; FDR = 5.23 × 10^-7; n = 36), aerobic electron transport chain (GO:0019646; FDR = 3.87 × 10^-6; n = 14), mitochondrial ATP synthesis coupled electron transport (GO:0042775; FDR = 3.87 × 10^-6; n = 14), purine nucleotide biosynthetic process (GO:0006164; FDR = 6.80 × 10^-6; n = 21), and electron transport chain (GO:0022900; FDR = 9.52 × 10^-6; n = 14).In **(B)** and **(C)**, bar length indicates enrichment significance as -log10(FDR).

Conserved genes were enriched for core housekeeping processes, including RNA metabolism, ribosome biogenesis, gene expression, ribonucleoprotein complex biogenesis, biosynthesis, and RNA processing (Fig. 2B). These enrichments indicate that genes with conserved regulatory architecture are concentrated in essential cellular programs related to RNA production, ribosome assembly, and general biosynthetic capacity.

In contrast, trans-only genes showed a distinct functional signature dominated by mitochondrial and respiratory processes (Fig. 2C). Enriched terms included mitochondrial translation, mitochondrial gene expression, cellular respiration, electron transport, oxidative phosphorylation-related processes, and nucleotide biosynthesis. These results suggest that trans-regulatory divergence is not randomly distributed across the genome but is concentrated in coordinated mitochondrial and energy-metabolism modules.

### Compensatory changes stabilize gene expression

Given the strong bias toward compensatory divergence, we investigated whether compensatory and reinforcing changes differ in the magnitude of their underlying regulatory components. If stabilizing selection permits larger regulatory mutations when they are offset by opposing changes, compensatory genes should exhibit larger absolute cis or trans effects than reinforcing genes (Metzger et al. 2017). To test this, we compared the absolute log2 fold-change magnitudes of net parental divergence, cis effects, and trans effects across regulatory classes (Fig. 3).

**Figure 3.**
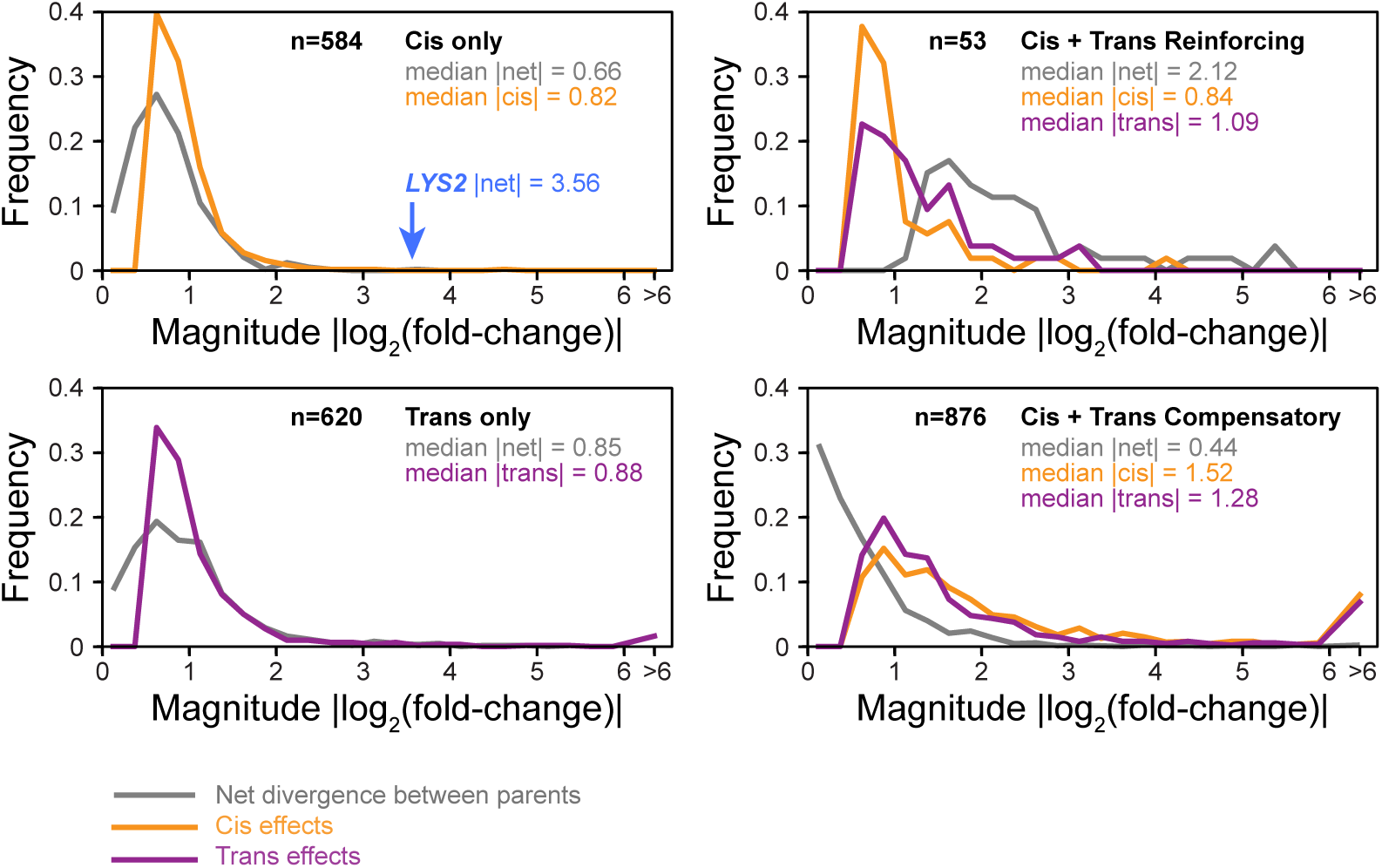
Compensatory cis–trans divergence permits large underlying regulatory changes while stabilizing net expression differences. Each panel shows the distribution of effect-size magnitudes, plotted as frequency versus |log2 (fold-change)|, for genes assigned to one regulatory class. Gray curves show the magnitude of net divergence between parents (parent–parent expression fold change). Orange curves show the magnitude of cis effects (allelic imbalance in the F1 hybrid, reflecting allele-linked local regulatory differences in a shared trans environment). Purple curves show the magnitude of trans effects (differences attributable to the regulatory environment after accounting for cis). Panel headers report the number of genes (n) and the median magnitude for each component. Regulatory classes are defined as follows: cis-only genes show significant cis effects with no detectable trans component; trans-only genes show significant trans effects with no detectable cis component; cis+trans reinforcing genes show cis and trans effects in the same direction, increasing net divergence; and cis+trans compensatory genes show cis and trans effects in opposite directions, yielding small net divergence despite large component magnitudes. *LYS2* is highlighted (blue arrow) as an extreme example within the cis-only distribution. Values exceeding 6 are grouped in the rightmost “>6” bin.

Consistent with this expectation, compensatory genes showed larger underlying regulatory component magnitudes than reinforcing genes (Fig. 3A and 3B). For cis effects, the median absolute effect was 1.52 in compensatory genes and 0.84 in reinforcing genes, corresponding to approximately 2.9-fold and 1.8-fold differences, respectively. Thus, the typical cis effect was approximately 1.8 times larger in compensatory genes than in reinforcing genes on the log2 effect-size scale. Trans effects showed a similar but more modest pattern, with median absolute effects of 1.28 in compensatory genes and 1.09 in reinforcing genes, corresponding to approximately 2.4-fold and 2.1-fold differences, respectively. These differences were statistically significant, with Mann–Whitney U tests confirming larger cis and trans effects in compensatory genes than in reinforcing genes (cis effects: p = 4.1 × 10−11; trans effects: p = 0.016).

Importantly, these larger cis and trans components did not produce larger net parental expression divergence. Instead, compensatory genes had much lower net parental divergence than reinforcing genes, with median |LFCnet| = 0.44 for compensatory genes compared with 2.12 for reinforcing genes (Fig. 3A, and 3B). This pattern is consistent with large cis and trans effects acting in opposite directions and partially canceling each other, thereby buffering total mRNA output despite substantial underlying regulatory divergence. In contrast, reinforcing genes showed larger net parental divergence because cis and trans effects act in the same direction. For comparison, cis-only and trans-only genes showed closer agreement between their single regulatory component and net parental divergence, as expected when divergence is dominated by one regulatory mode (Fig. 3C, and 3D). Together, these results indicate that compensatory evolution in this system often involves large opposing cis and trans changes that stabilize gene expression at the level of total transcript abundance.

### Hybrid expression patterns differ across cis–trans regulatory architectures

After estimating cis and trans regulatory components, we compared total expression in the F1 hybrid with expression in the two parental species (Fig. 4A). Across 5,386 genes in the hybrid expression analysis, hybrid expression was predominantly conservative. A total of 4,393 genes (81.6%) were classified as nondifferential, meaning that hybrid expression was not significantly different from either parent. In contrast, 143 genes (2.7%) showed additive expression, 819 genes (15.2%) showed expression biased toward one parent, and only 31 genes (0.6%) had hybrid expression outside the parental range. Thus, even in the presence of widespread regulatory divergence, most hybrid transcripts remained within the range of parental expression.

**Figure 4.**
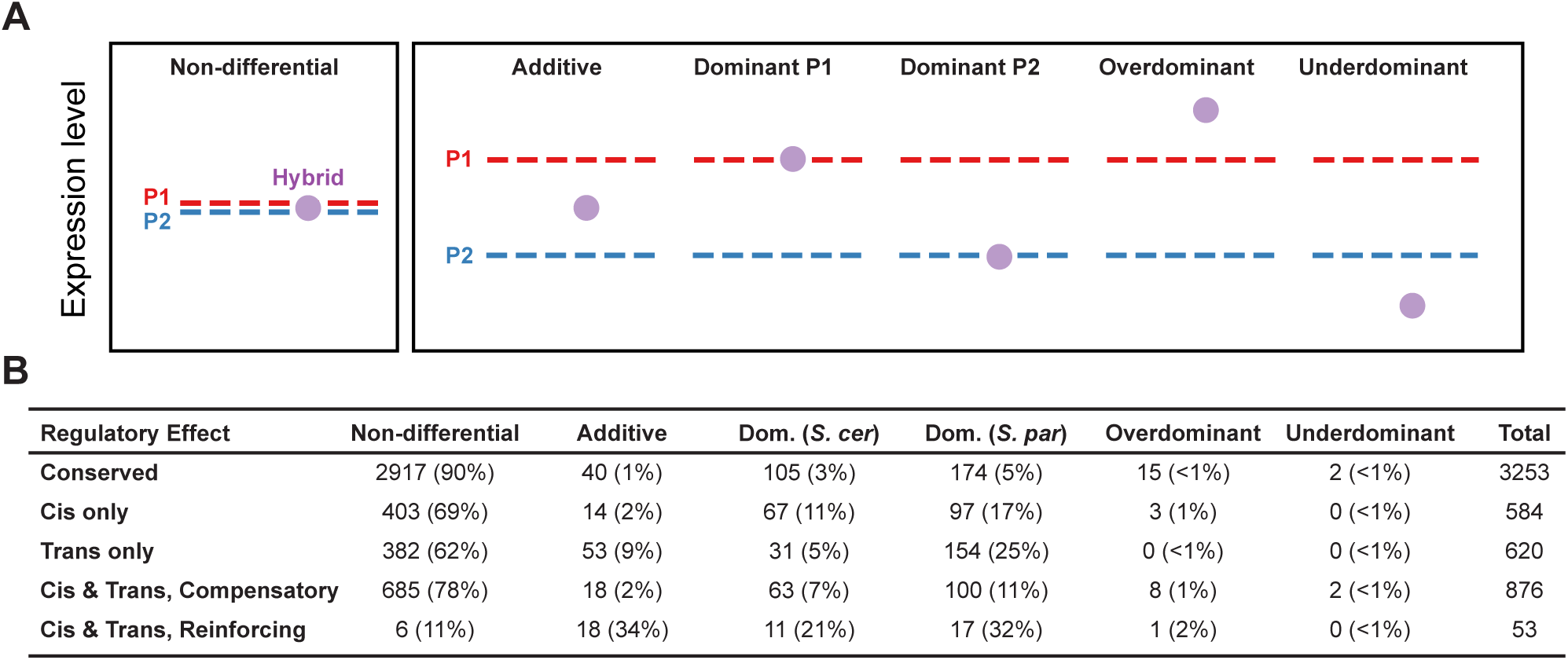
Hybrid expression modes and their distribution across regulatory classes. **(A)** Schematic of hybrid expression categories relative to the two parents. Hybrid expression was classified as non-differential when it fell within the parental range and was not significantly different from either parent; additive when it fell between the parents and differed from both; dominant when it matched one parent more closely than the other; and transgressive when it fell outside the parental range, either above both parents as overdominant or below both parents as underdominant. Purple circles denote hybrid expression. **(B)** Distribution of hybrid expression categories within each regulatory class. Values are shown as the number of genes, with percentages in parentheses, for non-differential, additive, dominance toward *S. cerevisiae*, dominance toward *S. paradoxus*, overdominant, and underdominant categories.

This conservative pattern was strongest for genes with no detectable cis or trans divergence. Among the 2,031 genes in the conserved regulatory class, 1,967 genes (96.8%) were nondifferential, as expected for genes without detectable regulatory divergence. Genes with only cis or only trans divergence were more likely to deviate from this pattern, but in different ways. In the cis-only class, 403 of 584 genes (69.0%) remained nondifferential, while the remaining genes were distributed mainly between *S. cerevisiae*-dominant and *S. paradoxus*-dominant expression. In the trans-only class, 382 of 620 genes (61.6%) were nondifferential, but genes showing parental bias were strongly skewed toward *S. paradoxus*. A total of 154 trans-only genes (24.8%) were classified as *S. paradoxus*-dominant, compared with only 31 genes (5.0%) classified as *S. cerevisiae*-dominant (Fig. 4B). Because trans-only genes were enriched for mitochondrial and respiratory functions in the GO analysis above, this dominance bias may reflect a coordinated trans-regulatory effect on mitochondrial and energy-metabolism pathways.

The clearest contrast emerged among genes carrying both cis and trans divergence. Most compensatory genes remained buffered, with 685 of 876 genes (78.2%) classified as nondifferential and another 18 genes (2.1%) classified as additive. Only a minority of compensatory genes showed dominant or transgressive expression patterns. By contrast, reinforcing genes were much less likely to remain nondifferential. Only 6 of 53 reinforcing genes (11.3%) were nondifferential, whereas 18 genes (34.0%) were additive, 11 genes (20.8%) were *S. cerevisiae*-dominant, and 17 genes (32.1%) were *S. paradoxus*-dominant (Fig. 4B). Therefore, opposite-sign cis-trans changes are usually associated with maintenance of hybrid expression within the parental range, whereas same-direction cis-trans changes are more likely to shift hybrid expression toward one parental state. This extends the previous result that compensatory genes can harbor large underlying regulatory differences while still producing stable total transcript output.

This comparison also helps explain the rare cases of transgressive expression, where hybrid expression falls outside the parental range. Reinforcing genes contributed to this small set of extreme expression patterns, but transgressive genes were more strongly enriched in the compensatory class. Among the 19 genes with transgressive expression and a resolved regulatory architecture, 10 belonged to the compensatory class (Fig. 4B). Although most compensatory genes remained well buffered, these rare failures of buffering were therefore disproportionately drawn from this class. Specifically, compensatory genes were 4.21-fold more likely than non-compensatory resolved genes to show overdominant or underdominant expression in the hybrid (Fisher’s exact test, p = 0.0023). Together, these results indicate that compensatory evolution usually stabilizes hybrid expression but can also create fragile regulatory equilibria in which mismatched or nonlinear cis-trans combinations occasionally push transcript levels beyond the parental range.

### Intron retention is highly conserved between *S. cerevisiae* and *S. paradoxus*

Introns are rare in budding yeast, occurring in only about 4-5% of genes, with only a few hundred introns across the genome (Spingola et al. 1999; Juneau et al. 2009). However, a much higher percentage of transcripts are spliced since some intron containing genes such as ribosome protein genes are highly expressed. Having shown that total gene expression in the hybrid is largely buffered despite widespread cis–trans divergence, we next asked whether post-transcriptional regulation showed a similar pattern of robustness. Allele-resolved intron-retention analysis revealed that most introns had low retention in both diploid parents and clustered near the diagonal in the parent–parent comparison (Fig. 5B), indicating broadly similar splicing efficiency between *S. cerevisiae* and *S. paradoxus* under these conditions. A small number of introns showed baseline species differences, including higher retention of *SFT1* in the *S. cerevisiae* parent and higher retention of *GPI15* in the *S. paradoxus* parent, but these differences were locus-specific rather than genome-wide.

**Figure 5.**
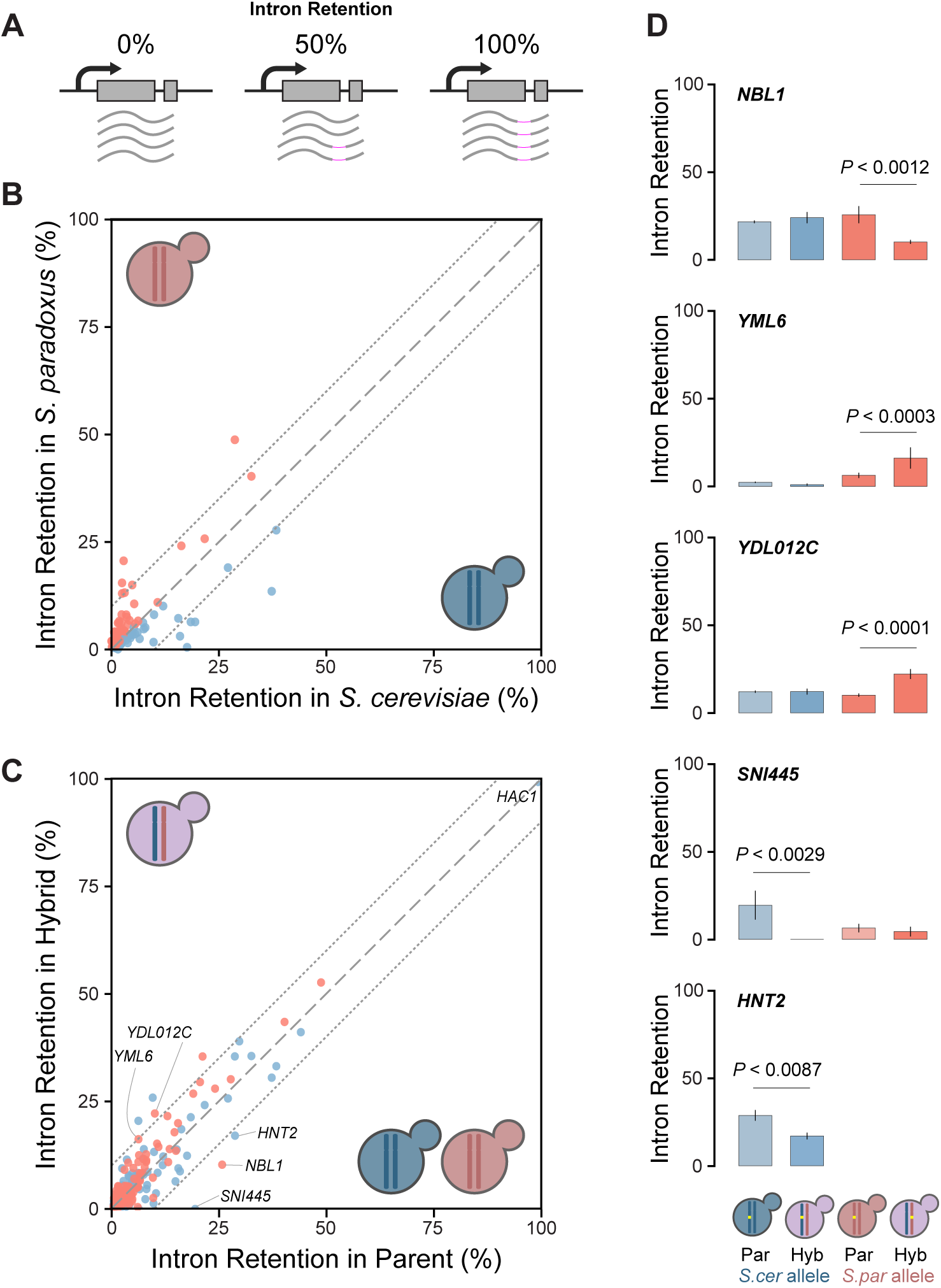
Intron retention is broadly conserved between *S. cerevisiae*, *S. paradoxus*, and their F1 hybrid, with rare allele-specific shifts. **(A)** Schematic of intron retention (IR) at 0%, 50%, and 100%. Gray arcs represent spliced reads, and magenta segments represent unspliced intron-containing reads. In panels **B–D**, blue denotes *S.c* or the *S.c* allele, and red denotes *S.p* or the *S.p* allele. Dashed lines in scatterplots indicate equal IR between axes, with flanking reference bounds where shown. **(B)** Parent-versus-parent comparison of mean IR. Each point represents one gene, summarized by the maximum mean IR among analyzed introns. Labeled genes are representative examples highlighting loci with higher retention in one parent or notable retention patterns. For example, *SFT1* showed higher retention in *S.c* than *S.p* (IR = 0.373 vs. 0.135; ΔIR = −0.238), whereas *GPI15* showed higher retention in *S.p* than *S.c* (IR = 0.488 vs. 0.287; ΔIR = +0.201). **(C)** Parent-versus-hybrid comparison of allele-resolved IR. Each point represents one analyzed intron comparing the parent with the corresponding hybrid allele. Labeled genes are representative examples of stable introns, modest shifts, or candidate hybrid-associated changes. Strong retention (IR ≥ 0.5) was rare across all comparisons. *HAC1* served as a retained-intron positive control and showed near-complete retention in the *S.c* parent and hybrid *S.c* allele (0.99 ± 0.000 vs. 0.98 ± 0.002), with no significant parent-to-hybrid shift (ΔIR = −0.0043; FDR = 0.175). **(D)** Representative hybrid-associated allele-specific IR shifts. Bars show mean IR ± SEM for parent (Par) and hybrid allele (Hyb). On the *S.p* allele, IR increased for *YDL012C* (0.10 ± 0.00 to 0.22 ± 0.02; ΔIR = +0.120; FDR = 1.24 × 10−5), decreased for *NBL1* (0.258 ± 0.049 to 0.102 ± 0.011; ΔIR = −0.155; FDR = 0.0077), and increased for near-threshold *YML6* (0.063 ± 0.015 to 0.162 ± 0.060; ΔIR = +0.099; FDR = 0.0029). On the *S.c* allele, IR decreased for *SNI445* (0.195 ± 0.082 to 0.000 ± 0.000; ΔIR = −0.195; FDR = 0.0157) and *HNT2* (0.288 ± 0.030 to 0.170 ± 0.019; ΔIR = −0.118; FDR = 0.0310). Horizontal brackets indicate the specific parent-to-hybrid allele comparison tested for each locus, with exact p values shown in the figure.

We then asked whether hybridization altered IR for each allele relative to its parental background. Parent-versus-hybrid comparisons again showed that most introns remained close to the diagonal (Fig. 5C), consistent with broadly stable splicing in the hybrid. Using an IR ratio greater than 0.1 as a descriptive threshold for retained introns, 22 of 178 introns (12.4%) were retained in both the *S. cerevisiae* parent and the hybrid *S. cerevisiae* allele. In the *S. paradoxus* comparison, retained introns increased modestly from 17 of 149 introns (11.4%) in the parent to 25 of 149 introns (16.8%) in the hybrid *S. paradoxus* allele. Strong intron retention, defined as IR ratio greater than or equal to 0.5, was uncommon and was driven almost entirely by the canonical retained intron in *HAC1*, which showed near-complete retention in both the *S.c* parent and the hybrid *S.c* allele. *HAC1* therefore served as positive control and showed no significant parent-to-hybrid shift. No introns in the *S.p* parent exceeded IR ≥ 0.5, whereas a single intron in the hybrid *S.p* allele, *GPI15*, exceeded this threshold. Thus, strong retention was rare and did not reflect a genome-wide hybrid effect. Together, these distributions indicate that hybridization does not broadly disrupt splicing efficiency, although the *S.p* allele set shows a modest right-shift in IR values.

Against this globally stable background, only a small number of introns showed clear hybrid-associated changes. Using an effect-size cutoff of |ΔIR| > 0.1 and FDR < 0.05, we identified four primary differential introns, plus one near-threshold event with strong statistical support (Fig. 5D). On the *S. paradoxus* allele, intron retention increased for *YDL012C* and decreased for *NBL1*, while *YML6* showed a near-threshold increase. On the *S. cerevisiae* allele, intron retention decreased for *SNI445* and *HNT2*. In each case, the opposite species allele showed little or no comparable shift, indicating that these responses are predominantly allele-specific rather than shared genome-wide effects of the hybrid background. Additional statistically significant but smaller shifts were detected below the effect-size cutoff, but these were treated as secondary events because their effect sizes were modest.

Together, these results show that post-transcriptional regulation is more stable than transcriptional regulation in this interspecific hybrid. Whereas total gene expression shows widespread cis–trans divergence across thousands of genes, intron retention is broadly conserved and only rarely reveals hybrid-associated, allele-specific shifts. This contrast suggests that the hybrid transcriptome contains extensive regulatory divergence at the level of expression output, but comparatively little disruption of splicing fidelity

### Promoter architecture underlies cis-regulatory divergence at *LYS2*

We examined the strongly cis-diverged gene *LYS2* as a case study for how promoter evolution may generate species-specific regulation (Fig. 3). The *S.c* promoter contains a 9-bp AT-rich insertion, TATGTATGA, located near −390 bp relative to the start codon (Fig. 6B). Because A/T-rich DNA is intrinsically less favorable for nucleosome formation, this insertion suggested that cis divergence at *LYS2* might reflect altered promoter architecture and local chromatin organization rather than simple gain or loss of a canonical activator-binding site.

**Figure 6.**
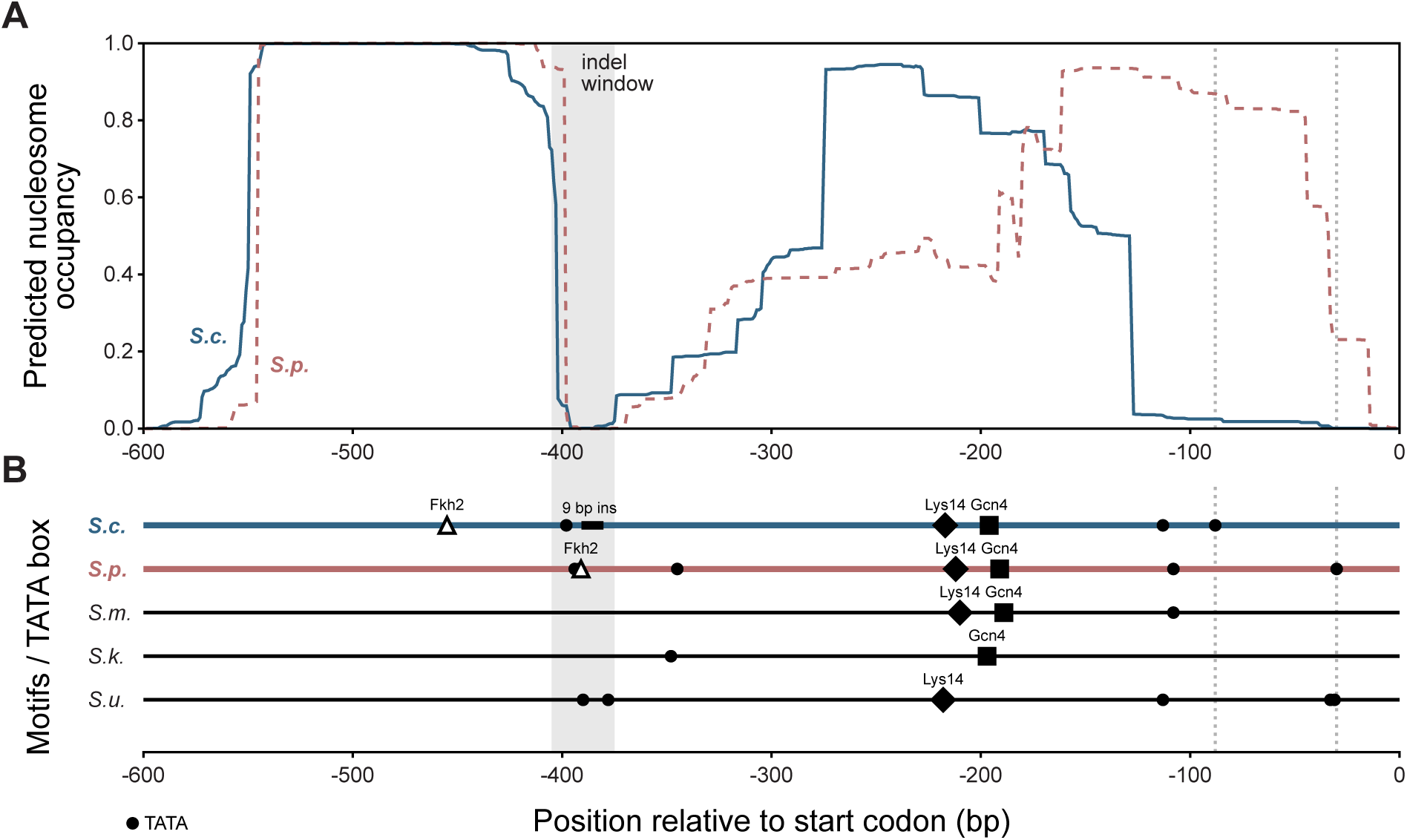
*LYS2* promoter architecture and predicted intrinsic nucleosome occupancy in *S. cerevisiae* and *S. paradoxus*. **(A)** NuPoP-predicted nucleosome occupancy across the 600 bp upstream of the *LYS2* start codon in *S. cerevisiae* (blue solid line) and *S. paradoxus* (red dashed line). The shaded region marks the indel-centered window (-405 to -375 relative to the start codon). Vertical dotted lines indicate promoter-proximal downstream positions highlighted in the comparative motif analysis. **(B)** Positions of candidate cis-regulatory features mapped onto the same coordinate system for *LYS2* promoters from *S. cerevisiae*, *S. paradoxus*, *S. mikatae*, *S. kudriavzevii*, and *S. uvarum*. Circles mark TATA-like sequence matches, triangles mark Fkh2-like matches, diamonds mark Lys14 sites, and squares mark Gcn4 sites. The shaded region marks the *S. cerevisiae*-specific 9-bp AT-rich insertion near -390.

Motif analysis supported this interpretation. Both species retained a conserved canonical Gcn4 site with the same match sequence, TGTGACTCAAT, at similar promoter positions, −196 bp in *S.c* and −191 bp in *S.p*. This argues against wholesale turnover of the major lysine-pathway regulatory input. In contrast, the best Fkh2-like candidate differed in both sequence and position. In *S.c*, the best match occurred at −455 bp with the sequence GTAAACA, whereas in *S.p*, the best match occurred at −391 bp with the sequence ATAAACA. TATA-like scanning further highlighted a species-specific architectural difference. Both promoters contained an upstream TATA-like sequence near the indel-rich region, TATATATA at −398 bp in *S.c* and TATATAAA at −394 bp in *S.p*, but only *S.c* carried an additional promoter-proximal TATA-like element at −88 bp, TATAAAAA (Fig. 6B). Thus, the key difference at *LYS2* is not the appearance of a new strong upstream activator motif, but a reorganization of promoter features in space.

These sequence changes produced marked differences in spacing among candidate regulatory elements. In *S.c*, the best Fkh2-like candidate was located approximately 57 bp upstream of the upstream TATA-like element. In *S.p*, the corresponding features were separated by only approximately 3 bp. Together with the additional downstream TATA-like element in *S.c*, these spacing shifts support a promoter-evolution model in which cis divergence arises through altered arrangement and phasing of promoter features rather than binary motif gain or loss.

Sequence-based nucleosome prediction reinforced this view. Predicted nucleosome occupancy is reported on a 0-to-1 scale, where values near 0 indicate low predicted occupancy and values near 1 indicate high predicted occupancy. Around the upstream insertion itself, mean predicted occupancy in the −405 to −375 bp window was low in both species, but lower in *S.c* than in *S.p* (0.078 versus 0.213). By contrast, predicted divergence in the core promoter was much stronger. In the −150 to −50 bp window, predicted occupancy was 0.132 in *S.c* compared with 0.882 in *S.p*. In the −120 to −80 bp window, predicted occupancy was 0.027 in *S.c* compared with 0.884 in *S.p*. These values predict a much broader promoter-proximal nucleosome-depleted region in *S.c*, coincident with the species-specific downstream TATA-like element.

An important control was that *LYS2* was not unusually diverged overall at the sequence level. Relative to 20 conserved-expression control promoters, *LYS2* showed high gap-free identity, 0.842, corresponding to approximately the 90th percentile among controls, but only typical indel burden, 34 indel bp, approximately the 60th percentile, and typical indel event count, 14 events, approximately the 65th percentile. Global A/T content was also similar between species, 0.617 in *S.c* and 0.615 in *S.p*, indicating that the relevant difference is not broad promoter degeneration or an overall base-composition shift. Instead, *LYS2* stood out specifically for local promoter architecture. Across the 20 conserved-expression controls, TATAWAWA matches were found only in far-upstream regions, −599 to −337 bp, and were absent from the promoter-proximal windows, −150 to −50 bp and −120 to −80 bp, in both species. In contrast, *LYS2* uniquely carried a proximal TATA-like element at −88 bp in *S.c*. Likewise, predicted promoter-proximal nucleosome divergence at *LYS2* was more extreme than in any control promoter. For the difference in predicted occupancy between *S.p* and *S.c*, the control maxima were 0.604 in the −150 to −50 bp window and 0.769 in the −120 to −80 bp window, whereas *LYS2* reached 0.750 and 0.857, respectively. By contrast, the difference around the upstream indel region was only 0.135 and remained within the control range. Thus, *LYS2* is not an outlier for overall promoter divergence, but it is an outlier for predicted promoter-proximal chromatin divergence.

Extending the comparison across additional sensu stricto species strengthened the same conclusion. TATA-like sequences near the upstream AT-rich region occurred in several species, but only *S.c* carried the promoter-proximal TATA-like element at −88 bp. *S. uvarum* had a downstream TATA-like feature near −33 bp, and *S.p* contained a short TATAAA sequence near −30 bp, but neither reproduced the *S.c* promoter configuration (Fig. 6B). Predicted promoter-proximal nucleosome occupancy followed these architectural differences. In the −150 to −50 bp and −120 to −80 bp windows, *S.c* remained low, 0.132 and 0.027, whereas *S.p* was high, 0.882 and 0.884. *S. kudriavzevii* and *S. uvarum* showed intermediate values, 0.462 and 0.401 for *S. kudriavzevii* and 0.551 and 0.480 for *S. uvarum*. *S. mikatae* showed near-maximal predicted occupancy, 0.997 and 0.999. Although the downstream TATA-like feature in *S. uvarum* was associated with a narrow local occupancy dip near −33 bp, 0.006 compared with 0.291 in *S.p*, it did not reproduce the broad promoter-proximal depletion predicted in *S.c* (Sup Fig. 1).

## DISCUSSION

Across the orthologous genes analyzed, regulatory divergence between *S.c* and *S.p* was common, but hybrid transcript levels were usually conservative. In the regulatory-class subset shown in Fig. 4B, 99.5% of genes remained within the parental expression range, whereas only 0.5% showed transgressive expression. For genes with both cis and trans contributions, compensatory cases exceeded reinforcing cases by approximately 16.5-fold. Thus, regulatory divergence frequently accumulated without corresponding large shifts in total transcript abundance, consistent with compensatory evolution. This interpretation agrees with hybrid-based and comparative studies that have repeatedly identified compensatory or antagonistic cis-trans divergence as a common feature of regulatory evolution (Landry et al. 2005; Tirosh et al. 2009; Metzger et al. 2017; Hill et al. 2021; Signor and Nuzhdin 2018). Together, these results support a model in which regulatory divergence is widespread but often cryptic at the level of total mRNA abundance because opposing cis and trans effects buffer transcript output.

The magnitude of the underlying regulatory effects further supports this buffering model. Metzger et al. (2017) found across *Saccharomyces* comparisons that compensatory genes tend to carry larger cis and trans effect sizes than reinforcing genes, consistent with stabilizing selection acting on total transcript output while allowing the underlying regulatory mechanisms to diverge. Our data fit that same qualitative expectation. However, F1 hybrid analysis measures the net cis and trans effects for each gene rather than identifying the individual mutations that produced them (Wittkopp and Kalay 2012; Hill et al. 2021). We therefore interpret the excess of compensatory effects as evidence for buffering at the level of transcript abundance, while remaining cautious about the exact mutational path at any individual locus.

Having established that compensatory divergence can buffer parental expression differences, the hybrid expression analysis allowed us to ask whether this buffering persists in the hybrid. Cis-only genes remained mostly within the parental range, but when they deviated they more often did so through additive or parent-biased outcomes. By contrast, trans-only genes were more strongly skewed toward dominance, particularly toward the *S.p* state. Similar relationships between cis divergence and additive inheritance, and between trans divergence and dominant or recessive inheritance, have been described in yeast and *Drosophila* hybrids (Schaefke et al. 2013; McManus et al. 2010; Wittkopp et al. 2004).

Our GO results further suggest that the trans component in this cross is not randomly distributed across genes (Fig. 4B, Fig. 2C). Trans-only genes were enriched for mitochondrial translation and respiratory functions, suggesting that coordinated trans-regulatory shifts may affect energy metabolism. Separating the trans-only class by hybrid expression category clarified this pattern further. The *S.p*-dominant trans-only genes carried a strong mitochondrial and respiratory signature, whereas the smaller *S.c*-dominant subset showed weaker functional enrichment. Our results suggest that there are one or more dominant trans-acting factors in *S. paradoxus* that regulate the expression of mitochondrial genes. This result agrees with yeast eQTL studies showing that trans-acting loci and hotspots can coordinately influence large gene modules (Brem et al. 2002; Yvert et al. 2003; Curtis et al. 2013; Albert et al. 2018).

At the same time, compensation in our data was associated with both robustness and fragility. Although most compensatory genes were buffered, 10 of the 19 transgressive genes with resolved regulatory architecture belonged to the compensatory class. Thus, compensatory cis-trans divergence usually helped maintain hybrid expression within the parental range, but in a small number of cases these opposing regulatory effects were associated with expression outside the parental range. This pattern is compatible with earlier work showing that antagonistic or novel cis-trans interactions can contribute to hybrid misexpression (Landry et al. 2005; Tirosh et al. 2009; McManus et al. 2010; Herbst et al. 2017). An important next question is how much additional misregulation will appear once meiotic recombination reshuffles these co-evolved regulatory components. In the F1, each allele remains embedded in an intact parental chromosome and experiences a trans environment that still contains both parental regulatory systems. Recombinant segregants should expose more non-coevolved combinations of cis elements and trans-acting alleles, and yeast segregant studies indicate that trans-regulatory architectures are often multilocus and modular (Albert et al. 2018). We therefore expect recombinant hybrid genomes to reveal a broader spectrum of transgressive expression and regulatory incompatibility than is visible in the F1 itself.

Post-transcriptional regulation provides a complementary perspective. Although only a small fraction of *Saccharomyces* genes contain introns, intron retention in budding yeast can be regulated and functionally important rather than simply reflecting splicing noise. Canonical examples include *HAC1*, whose intron controls translation during the unfolded protein response, and *YRA1*, whose intron participates in autoregulation of mRNA export factor abundance (Cox and Walter 1996; Ruegsegger et al. 2001; Rodriguez-Navarro et al. 2002; Preker and Guthrie 2006). More broadly, introns have been implicated in starvation responses and dynamic physiological programs in yeast, and work across eukaryotes has established intron retention as a regulated mode of gene control (Gonzalez-Hilarion et al. 2016; Jacob and Smith 2017; Parenteau et al. 2019; Gomez-Montalvo et al. 2024). In contrast to the widespread transcriptional divergence observed here, allele-resolved intron retention was remarkably conserved. Most introns showed low retention in both parents, with the exception of *HAC1*, which shows high intron retention, as expected. Only a handful of loci showed clear hybrid-associated allele-specific shifts in intron retention. GO enrichment of the intron-containing genes shows that these genes are involved core gene-expression and ribosome-associated pathways including translation, protein biosynthesis, ribosome biogenesis, rRNA processing, and ribonucleoprotein-complex biogenesis. These same GO terms are enriched in genes whose expression is conserved between *S. cerevisiae* and *S. paradoxus* (Fig. 2B). The observation that intron-containing genes are affect core biological processes and show high conservation in gene expression levels between species could explain why we find only a few quantitative differences in intron retention between species and in the hybrid.

Finally, while the genome-wide analyses reveal the scale and consequences of regulatory divergence, they do not identify the sequence features that generate strong cis effects at individual loci. Cis-regulatory divergence often cannot be reduced to the gain or loss of a single obvious transcription factor binding site. In yeast, promoter function is closely related to local chromatin context, nucleosome organization, and sequence composition, all of which can influence transcriptional responsiveness and evolutionary change (Basehoar et al. 2004; Tirosh and Barkai 2008; Hornung et al. 2012; Rosin et al. 2012). Sequence features such as AT-rich tracts can disfavor nucleosome formation and contribute to local chromatin structure (Segal et al. 2006; Kaplan et al. 2009; Segal and Widom 2009). Locus-level analyses of promoter architecture and sequence-based nucleosome prediction can therefore provide useful mechanistic context for strong cis effects, even when direct chromatin measurements are not available (Xi et al. 2010). Promoter sequence, therefore, is not merely correlated with regulatory divergence; it can help explain how that divergence arises mechanistically.

The *LYS2* case study provides a mechanistic foothold for one strong cis-diverged locus. The most informative differences were not wholesale turnover of the canonical Gcn4 input, which was conserved in both species, but local promoter architecture, an *S.c*-specific AT-rich insertion, a repositioned Fkh2-like candidate motif, altered spacing among upstream features, and a promoter-proximal TATA-like element found only in *S.c*. This type of local reorganization is fully consistent with current views of cis-regulatory evolution, in which small substitutions and indels alter binding, spacing, and cooperative context rather than simply creating or destroying one master site (Wray 2007; Wittkopp and Kalay 2012; Shih and Fay 2021). In yeast, TATA-containing promoters define a distinct regulatory class, promoter nucleosome architecture is closely linked to transcriptional plasticity, and promoter sequence can encode substantial differences in intrinsic nucleosome preference (Basehoar et al. 2004; Tirosh and Barkai 2008; Segal et al. 2006; Kaplan et al. 2009; Hornung et al. 2012; Rosin et al. 2012; Xi et al. 2010). Within that framework, the *LYS2* cis effect is directionally predictable. The *S.c* promoter is predicted to support a broader promoter-proximal nucleosome-depleted region than the *S.p* promoter, and that architectural difference provides the most parsimonious explanation for the allele-specific expression difference we observe. Because *LYS2* is not an outlier for overall promoter divergence but is an outlier for predicted proximal chromatin divergence, our results specifically implicate promoter architecture and intrinsic chromatin organization rather than generalized promoter decay.

These mechanistic interpretations should be viewed within several boundaries of inference. We measured expression and intron retention in one growth condition and one F1 hybrid combination, and RNA-level buffering does not necessarily imply equivalent protein output. Even with those caveats, the broader picture is clear. Between *S.c* and *S.p*, regulatory divergence is widespread but often hidden because opposing cis and trans changes stabilize total expression. By contrast, intron retention remains largely conserved, and *LYS2* illustrates how a strong cis effect can emerge through localized changes in promoter architecture and predicted chromatin organization. The interspecific hybrid therefore appears less like a transcriptomic failure state than a buffered intermediate, stable in overall output, but carrying cryptic regulatory divergence that may become more visible when parental alleles are reshuffled in recombinant segregants.

## DATA AVAILABILITY

The short-read sequencing data reported in this study have been deposited to the NCBI BioProject database, accession number PRJNA1469180.

## ACKNOWLEDGEMENTS

We thank Patricia Wittkopp, David Zapulla, and members of the Lang Lab for comments on the manuscript.

## FUNDING

This work was supported by a grant from the National Institutes of Health (R35GM149540). Portions of this research were conducted on Lehigh University’s Research Computing infrastructure partially supported by the National Science Foundation (Award 2019035).

## COMPETING INTERESTS

The authors declare no conflicts of interest.

## AUTHOR CONTRIBUTIONS

Conceptualization: GIL

Experiments: DR

Analysis: DR

Writing: DR, GIL

Editing: DR, GIL

Funding Acquisition: GIL

---

**Supplemental Figure 1.**
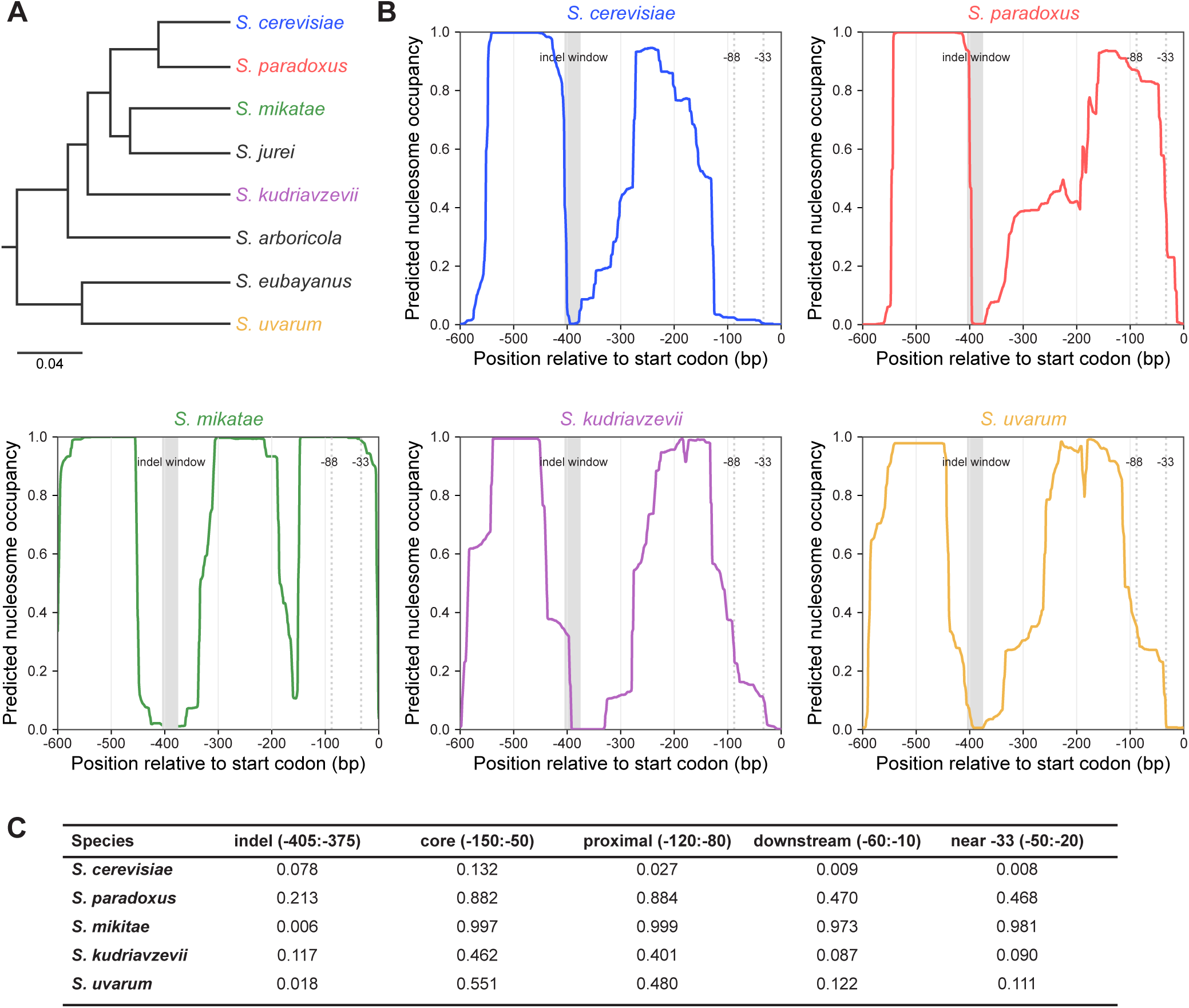
Multi-species comparison of *LYS2* promoter context and predicted intrinsic nucleosome occupancy across *Saccharomyces* sensu stricto. **(A)** Phylogenetic context for the Saccharomyces species included in the comparative *LYS2* promoter analysis; species analyzed for nucleosome occupancy are highlighted in color. **(B)** NuPoP-predicted intrinsic nucleosome occupancy across the 600 bp upstream region of *LYS2*, plotted relative to the ATG start codon for *S. cerevisiae*, *S. paradoxus, S. mikitae, S. kudriavzevii,* and *S. uvarum.* Occupancy values range from 0 to 1, with lower values indicating predicted nucleosome depletion and higher values indicating predicted nucleosome occupancy. The gray shaded region marks the indel-centered window (-405 to -375 bp), and dotted vertical lines mark the promoter-proximal positions at -88 and -33 bp. **(C)** Mean predicted occupancy in the indicated promoter windows. *S. cerevisiae* shows the strongest promoter-proximal depletion in the -150 to -50 bp and -120 to -80 bp windows, whereas *S. paradoxus* and *S. mikatae* show high predicted occupancy, with *S. kudriavzevii* and *S. uvarum* showing intermediate profiles.

## Notes

### Competing Interest Statement

The authors have declared no competing interest.

